# “Tiny Biome Tales”: a gamified review about the influence of lifestyle choices on the human microbiome

**DOI:** 10.1101/2024.06.29.601357

**Authors:** Matthias Schweitzer, Maximilian Wlasak, Birgit Wassermann, Florian Marcher, Christian Poglitsch, Johanna Pirker, Gabriele Berg

## Abstract

In the last two decades, new discoveries from microbiome research have changed our understanding of human health. It became evident that daily habits and lifestyle choices shape the human microbiome and ultimately determine health or disease. Therefore, we developed “Tiny Biome Tales” (https://microbiome.gamelabgraz.at/), a science pedagogy video game designed like a scientific review based exclusively on peer-reviewed articles, to teach about the influence of lifestyle choices on the human microbiome during pregnancy, early and adult life, and related health consequences. Despite the scientific character, it can be played by a broad audience. Here, we also present a scientific assessment, and showed that playing the game significantly contributed to knowledge gain. The innovative style of the “gamified review” represents an ideal platform to disseminate future findings from microbiome research by updating existing and adding new scenes to the game.

## Introduction

Microorganisms and their functions have widespread, significant, and generally positive impacts on human health ^1^. A relationship so vital that it can be viewed as one ecological unit called holobiont ^2^. Consequently, maintaining a balance in the relationship between microorganisms and their host is essential for promoting health in the state of eubiosis and for the onset of a disease in dysbiosis, which emphasizes the fact that all diseases correlate to the microbiome ^3^. Research suggests that certain aspects of the human microbiome are predetermined by host genetics ^4, 5^, geographical location ^6, 7^ and ethnicity ^8, 9^. However, these predetermined drivers of the human microbiome may be better explainable by lifestyle factors linked to specific geographical locations and ethnicity, including diet, hygiene, and culture ^10, 11^. Indeed, the human microbiome exhibits high individual variability ^12, 13^, a trait that is more pronounced in urban settings ^11^ and might further increase with predicted urbanization trends ^14^. Considering both, the inter-individual differences in the human microbiome, and the impact of several decision-based factors such as diet ^15, 16^, exercise ^17^, the frequent use of disinfectants and germicidal soaps ^18^, being vaccinated or treated for an infection ^19^, or having a companion dog ^20^, it becomes evident that, to certain degree, daily habits and lifestyle choices shape the human microbiome. Even prior to birth, maternal dietary habits were shown to have an impact on the microbiome of the infant ^21^. Mode of delivery and the choice between formula and breast-feeding influence the development of the child’s microbiome ^22^. The role of early-life microbiome development was particularly emphasized repeatedly ^23, 24^. One example is the described causal relationship between an immature microbiome and malnutrition ^25, 26^. As we continue to advance our knowledge of microbiomes, it becomes evident that microbiome research is integral for public health matters ^23^, especially by explaining potential causality and communicability of certain diseases traditionally considered noncommunicable ^24^. The paradigm of the relationship between microbes and disease is shifting towards microbiome and health. Functional diversity matters and is a key factor for a healthy microbiome and thus for a healthy host ^3^. Translation of microbiome research, in both educational and extracurricular settings, offers potential to influence individual lifestyle choices, and thereby prevention strategies for these chronic diseases ^27^.

Microbiome research has received attention in popular press over the past few years, where often the impact of lifestyle choices such as diet, exercise, and probiotics on microbiome-related health outcomes are highlighted ^29^. Consequentially, there is an increasing awareness of the term microbiome in society. Blanton et al. called for a proactive public dialogue about benefits and risks of microbiome-directed interventions for malnutrition in early-life, and emphasized the importance of educating about the microbiome and its impact on health. It is time for universities to adapt and enhance their educational approaches and public outreach creatively ^28^. Now seems to be the right time to educate the public, policymakers and public health practitioners about microbiome science ^23^.

Participatory action research was found to be effective in educating citizens about lifestyle choices, their impact on the gut microbiome, and potential links with noncommunicable diseases, as well as improving self-care towards a healthy gut microbiome, particularly among participants at higher risk of developing noncommunicable diseases ^30^. The integration of game elements within educational settings, commonly referred to as gamification ^31^, is another promising approach. This was shown in several science pedagogy board games such as “Gut Check: The Microbiome Game”, developed to teach about microbes in the human gut in a challenging and fun way, with a focus on beneficial microbes ^32^. An alternative approach is the one described for the game “Microbial Pursuit”. Students actively learn the subject while formulating questions and answers, creating the game themselves, making “Microbial Pursuit” an interesting tool for educational units including revision of the content ^33^. “BacteriaGame”, an educational card game about bacteriology was employed in a range of educational settings. Compared to a traditional lecture, playing the “BacteriaGame” resulted in a significantly higher learning outcome ^34^. Indeed, the implementation of active learning methods and gamification into education on science, technology, engineering, and mathematics (STEM) has demonstrated a positive influence on both student achievement and motivation ^35, 36^, also particularly benefiting disadvantaged students ^37^, but an interactive game that explains the enormous progress of microbiome research for our health is missing.

In this study, we present and evaluate “Tiny Biome Tales”, a science pedagogy video game developed to teach about the influence of lifestyle choices on the human microbiome during pregnancy, early life, and adult life, and the related health consequences. “Tiny Biome Tales” can be considered as “gamified review”, based exclusively on peer-reviewed articles, that was intended for a broad audience and designed to be playable by students and interested members of the public from adolescence onwards. For this purpose, we aimed to translate scientific terminology into language that is understandable by a broad audience but is still neutral and accurate. The quality of the game was evaluated using a questionnaire based on the MEEGA+ model ^38^, a tool designed to assess educational video games, which is a modified version of the widely used MEEGA questionnaire ^39^. Moreover, the effectiveness of the learning process was evaluated by comparing a group that completed knowledge assessments after playing the game to another group that completed the same assessments prior to gameplay. The game will be incorporated into the Massive Open Online Course (MOOC) “Microbiome & Health” that provides basic and applied knowledge about the microbiome (https://imoox.at/course/microbiome).

## Results

### “Tiny Biome Tales”: a gamified review about the influence of lifestyle choices on the human microbiome

The science pedagogy video game “Tiny Biome Tales” (https://microbiome.gamelabgraz.at/) was developed to provide a captivating and motivating learning experience about the human microbiome and related health consequences. Players can explore factors known to shape the human microbiome by selecting in-game elements (Fig. 1). With each choice, players are provided with concise information on the selected topic. Furthermore, choosing elements for the first time unlocks an entry in the so-called “Codex”. These codex-entries contain more extensive information on the corresponding content for individuals interested in learning more on certain topics. Each microbial species mentioned in the game is accompanied by its own codex entry, offering supplementary details about the species and its potential impact on human health. The players can track their progress by utilizing the “Diversity Index” element in the game. A higher Diversity Index indicates a greater number of correct decisions. The Diversity Index is a simplified metric rooted in foundational research suggesting that increased microbial diversity correlates with health ^3^. Although the indicators of a healthy microbiome are still under debate, functional diversity reflected in structural diversity as well is a key component that represents ecosystem resilience and health ^3^. It should be noted that the Diversity Index primarily serves as a gameplay element designed to enhance engagement rather than a strictly scientific outcome. We are encouraged to carefully consider the decisions, and the game aims to demonstrate the overall impact of the decisions as a whole. It emphasizes that there are not just right and wrong decisions, but rather a sum of their effects, represented by the Diversity Index. All information presented is provided with scientific references, each of which is documented in a codex entry with a DOI URL to peer-reviewed articles that is hyperlinked within the game. While aiming to translate scientific terminology into a language that is understandable by a broad audience, the resulting in-game text content still reads more like a scientific review article than what is expected from a video game. In conjunction with the scientific citation style, this has led to the development of an educational video game that may also be referred to as a “gamified review”. Currently, the game features nine interactive scenarios, some of which are multi-stage, that cover the impact of various lifestyle choices from pregnancy to old age (Fig. 2). In the game’s narrative, players begin as a pregnant character faced with decisions that impact both their own well-being and that of their unborn child. Decisions include nutrition, medication, childbirth method, and infant feeding. The game progresses through early years, allowing players to choose their playground environment, pet ownership, and food preferences. Adulthood brings opportunities for social events, exercise, and interpersonal relationships in a fun and engaging manner. The game concludes with old age, where players make decisions to encourage healthy aging. All information provided in “Tiny Biome Tales” is based on current scientific evidence, however, the game also represents a perfect platform to disseminate future findings from microbiome research through updating existing and adding new scenes to the game.

**Fig. 1:**
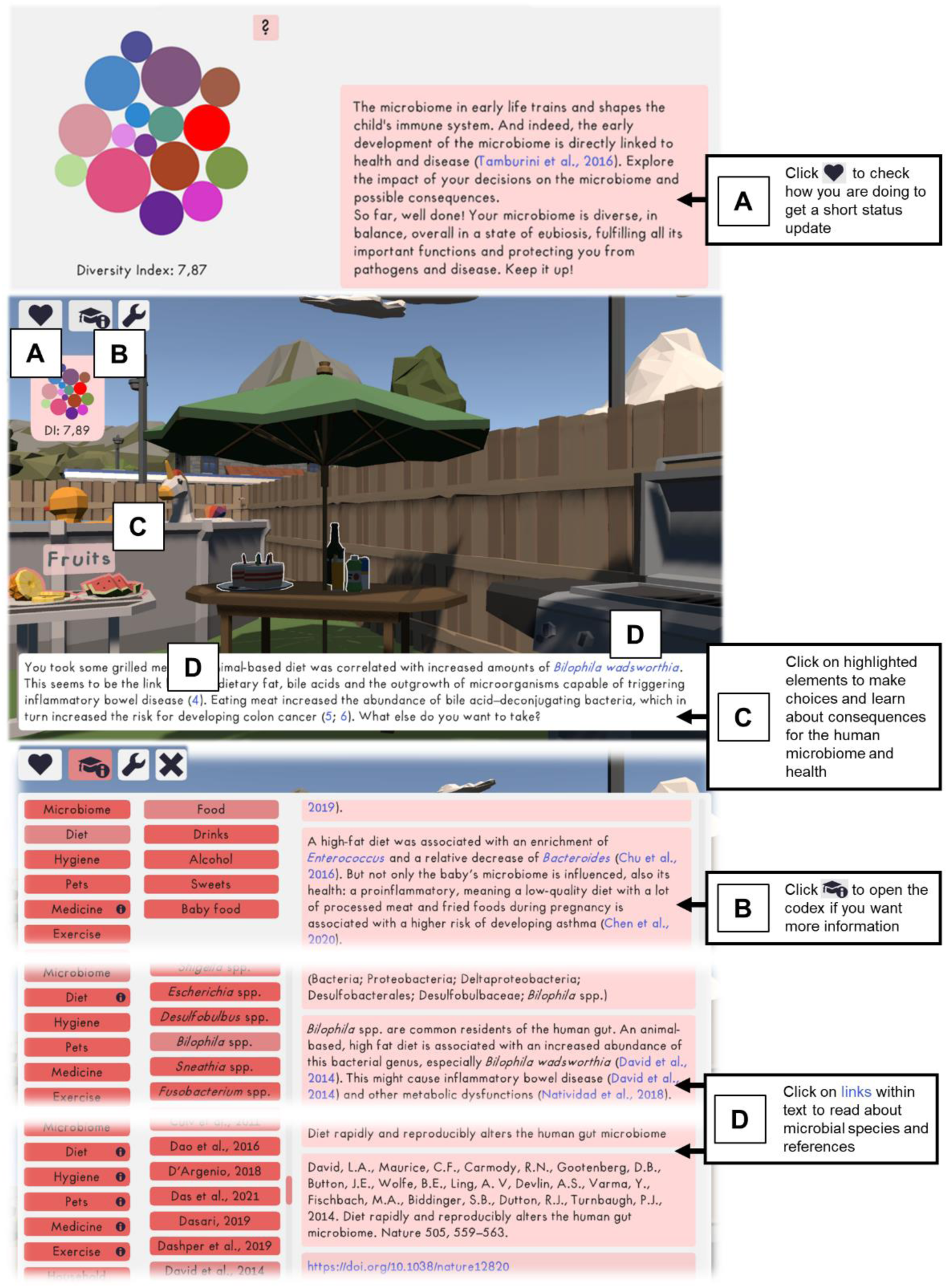
Visual excerpt of “Tiny Biome Tales” depicting options for lifestyle choices and health-related consequences thereof. The player makes lifestyle decision by clicking on highlighted elements, each lowering or increasing the “Diversity Index”, i.e., a simplified display of the playeŕs microbiome health status. Each decision made displays related information and unlocks entries in the so-called “Codex”, providing immediate insight into the impact of the decisions on the human microbiome and health, with additional information available for those who wish to delve deeper.

**Fig. 2:**
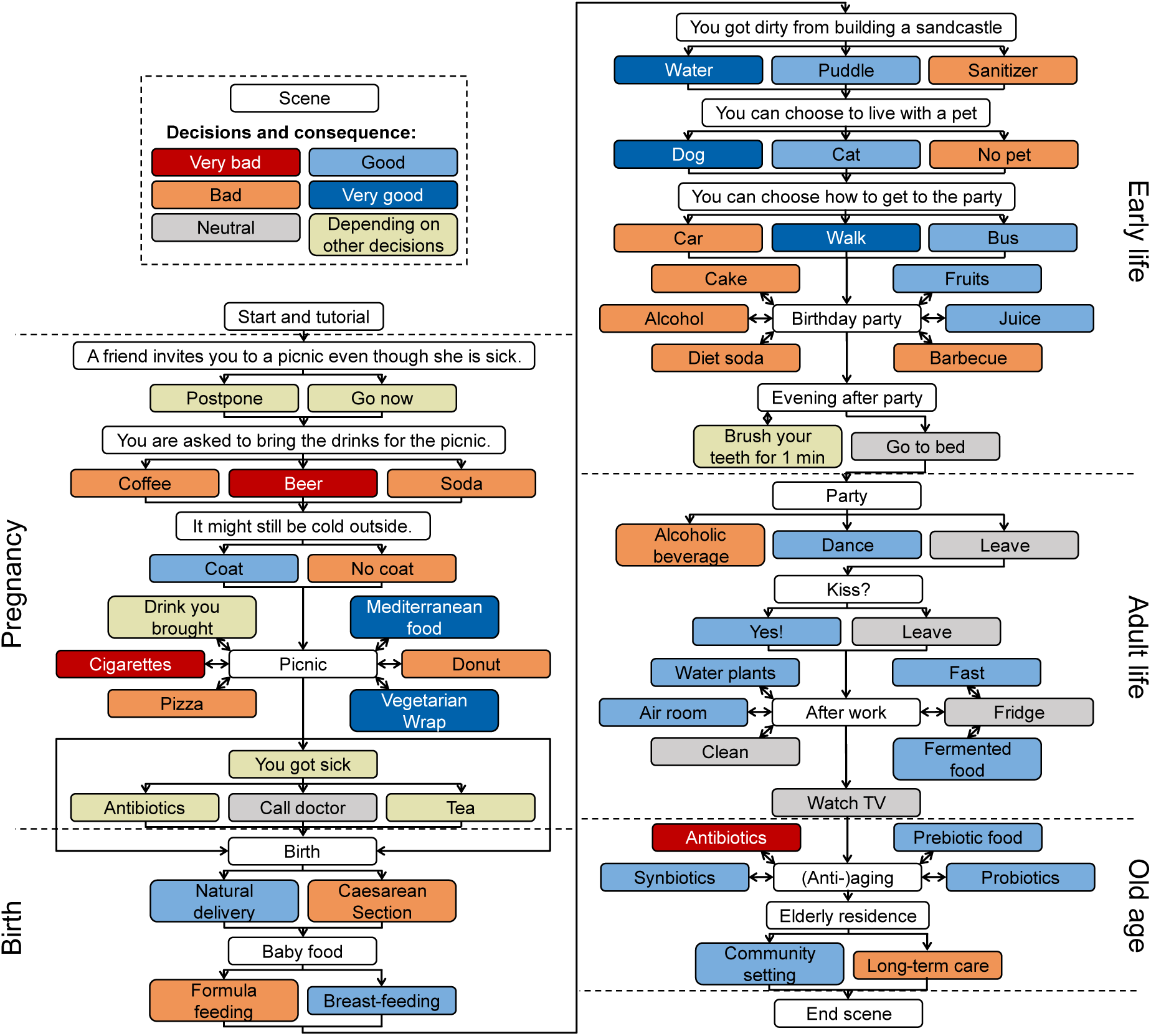
Summary of all scenes and associated options currently available for players of “Tiny Biome Tales”. The colors indicate positive (light to dark blue) or negative (orange to red) influence of each decision on the human microbiome and health.

### After playing “Tiny Biome Tales”, participants got significantly more knowledge questions right

A pilot study was performed to evaluate the quality of “Tiny Biome Tales” and the learning success of playing (Fig. 3 A). A total of 65 participants completed the questionnaires, categorized as either “experts” (*N* = 22), consisting of educators, students, and researchers specialized in microbiome research, or “non-experts” (*N* = 43). The non-expert group was further divided into a “control” group (*N* = 20) who completed a knowledge questionnaire before engaging with the game, and a “test” group (N = 23) who completed the same questionnaire after playing the game to measure the impact on learning outcomes. Demographic information of all participants was surveyed (Table 1). As anticipated, the expert group rated their knowledge of the human microbiome higher than the non-expert control and test groups (*P.adj.* < 0.001). The significantly younger (*P.adj.* < 0.001) control (*N* = 20) and test groups (*N* = 23) consisted mainly of 18–28-year-old students and individuals from the IT sector.

**Fig. 3:**
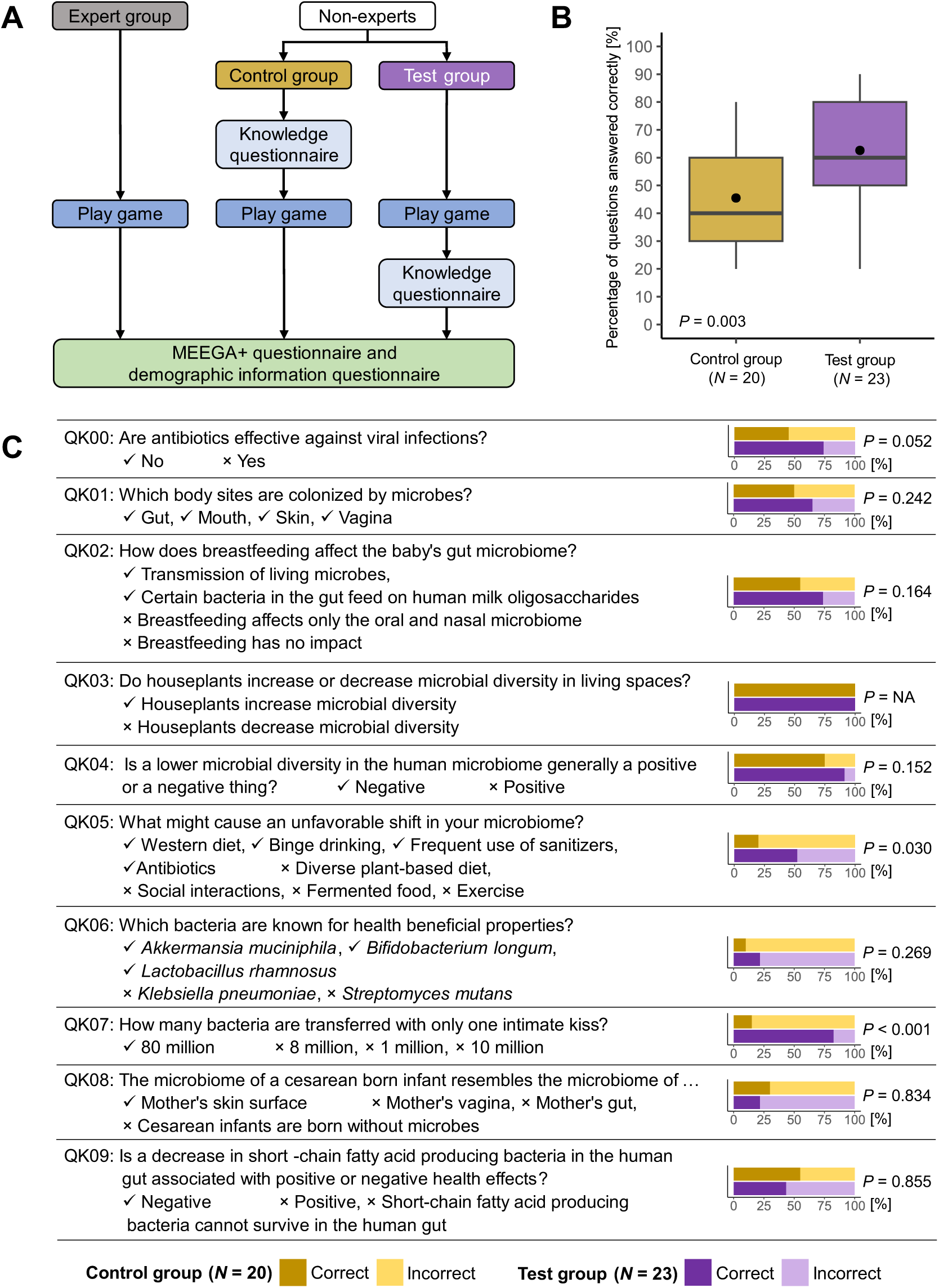
Study design and learning success with playing “Tiny Biome Tales”. **A** Design of the pilot study for evaluating the quality and learning outcome of the educational video game “Tiny Biome Tales”. **B** The percentage of questions answered correctly, displayed for both the control group (N = 20) and the test group (N = 23). One-sided Mann–Whitney *U* test was performed to assess if the test group significantly performed better than the control group. **C** The percentage of participants from both the control group (*N* = 20) and the test group (*N* = 23) who answered the knowledge questions correctly or incorrectly. One-sided isher’s exact tests were performed to assess if the test group significantly performed better than the control group.

**Table 1:** Demographic information of study participants.

| ID: Question | Control (N=20) | Test (N=23) | Expert (N=22) |
| --- | --- | --- | --- |
| QD00: What is your age group? |  |  |  |
| 18-28 | 19 (95.0%) | 23 (100.0%) | 6 (27.3%) |
| 29-39 | 1 (5.0%) | 0 (0.0%) | 13 (59.1%) |
| 40-50 | 0 (0.0%) | 0 (0.0%) | 3 (13.6%) |
| <sup>a</sup>$P < 0.001$ | a | a | b |
| QD01: What gender do you identify as? |  |  |  |
| Male | 10 (50.0%) | 15 (65.2%) | 10 (45.5%) |
| Female | 9 (45.0%) | 7 (30.4%) | 12 (54.5%) |
| Unspecified | 1 (5.0%) | 1 (4.3%) | 0 (0.0%) |
| <sup>b</sup>$P = 0.406$ | | | |
| QD02: What is your profession? |  |  |  |
| Student | 13 (65.0%) | 17 (73.9%) | 2 (9.1%) |
| Educational Worker | 1 (5.0%) | 0 (0.0%) | 10 (45.5%) |
| Researcher | 0 (0.0%) | 0 (0.0%) | 9 (40.9%) |
| IT | 4 (20.0%) | 3 (13.0%) | 0 (0.0%) |
| Business | 1 (5.0%) | 2 (8.7%) | 1 (4.5%) |
| Worker | 1 (5.0%) | 1 (4.3%) | 0 (0.0%) |
| <sup>b</sup>$P < 0.001$ | a | a | b |
| QD03: Do you consider yourself literate in the usage of computers? |  |  |  |
| Very much | 4 (20.0%) | 7 (30.4%) | 7 (31.8%) |
| Yes | 13 (65.0%) | 12 (52.2%) | 10 (45.5%) |
| A little | 3 (15.0%) | 3 (13.0%) | 5 (22.7%) |
| Not at all | 0 (0.0%) | 1 (4.3%) | 0 (0.0%) |
| <sup>a</sup>$P = 0.930$ | | | |
| QD04: How often do you play video games? |  |  |  |
| Daily: every day | 3 (15.0%) | 2 (8.7%) | 2 (9.1%) |
| Weekly: at least once a week | 7 (35.0%) | 5 (21.7%) | 4 (18.2%) |
| Monthly: at least once a month | 0 (0.0%) | 3 (13.0%) | 3 (13.6%) |
| Rarely: From time to time | 8 (40.0%) | 11 (47.8%) | 10 (45.5%) |
| Never | 2 (10.0%) | 2 (8.7%) | 3 (13.6%) |
| <sup>a</sup>$P = 0.605$ | | | |
| QD05: How much do you think you know about the human microbiome? |  |  |  |
| A lot | 0 (0.0%) | 0 (0.0%) | 9 (40.9%) |
| The basics | 8 (40.0%) | 9 (39.1%) | 13 (59.1%) |
| A little | 7 (35.0%) | 11 (47.8%) | 0 (0.0%) |
| Nothing | 5 (25.0%) | 3 (13.0%) | 0 (0.0%) |
| <sup>a</sup>$P < 0.001$ | a | a | b |

<sup>a</sup> Kruskal-Wallis test; <sup>b</sup> Fisher's exact test; The letter code indicates significant differences ( $P_{adj.} < 0.001$ ) between groups, determined via Dunn's post hoc test or post hoc pairwise Fisher's exact test and correcting $P$ -values using Benjamini-Hochberg.

With this preliminary study (Fig. 3 A), we were able to demonstrate the educational value of the game. Participants from the test group got on average 62.6% of the questions right after playing the game, significantly more compared to control group with an average of 45.5% (*P* = 0.003; Fig. 3 B). Overall, the test group outperformed the control group in seven out of ten questions (Fig. 3 C). All participants of both the control and the test group got Q 03 “Do houseplants increase or decrease microbial diversity in living spaces” right. Q 05 “What might cause an unfavorable shift in your microbiome?” was answered correctly by 52.2% of the test group participants, a significantly higher percentage compared to the 20% of the control group (*P* = 0.030). At Q 07 “How many bacteria are transferred with only one intimate kiss?” 82.6% of the test group selected the right answer “80 million” ^40^, compared to 15% of the control group (*P* < 0.001). Interestingly, at two questions a higher percentage of control group members answered correctly compared to the test group, although not significantly. QK08 was answered correctly by 30% of the control group and by 21.7% of the test group, and QK09 by 55% of the control group and 43.5% of the test group.

### “I would recommend this game to people willing to learn about the subject”

All study participants were asked to rate their player experience on a Likert scale from 1 (strongly disagree) to 5 (strongly agree) in a questionnaire based on the MEEGA+ model ^38^ (Fig. 4). At QE00A – “There was something interesting at the beginning of the game that captured my attention”, 77.3% of the expert group (arithmetic mean M on ikert scale of 4.0), 87.0% of the test group (M = 4.4) and even 100% of the control group (M = 4.3) agreed or strongly agreed. This outcome was exactly what we intended with the very first in-game scene, a bright screen with the words “You got pregnant” and a “Continue” button. However, only 41.3% of all players agreed that they lost track of time (QE00B - control group M = 3.5; test group M = 3.8; expert group M = 3.0) and 49.2% agreed that they forgot their immediate surroundings while playing (QE00C - control group M = 3.3; test group M = 3.7; expert group M = 3.1). Overall, 87.7% of the participants had fun during the game (QE01A - control group M = 4.1; test group M = 4.4; expert group M = 4.0) and 81.5% agreed that parts of the game made them smile (QE01B - control group M = 4.4; test group M = 4.2; expert group M = 4.0). There was considerable disagreement about the statement “The game is appropriately challenging for me”. 35. % of all study participants rated this statement with disagree or strongly disagree (QE02A - control group M = 3.1; test group M = 3.0; expert group M = 3.1). However, 66.2% of all players agreed that the game provides new challenges at an appropriate pace (QE02B - control group M = 3.8; test group M = 3.8; expert group M = 3.5) and that the game does not become monotonous (QE02C - control group M = 3.8; test group M = 4.0; expert group M = 3.4). The majority of the study participants were confident that the game would be easy for them when looking at the play screen for the first time (QE03A - control group M = 3.9; test group M = 3.7; expert group M = 4.1) and the game’s structure and content helped them to learn with the game (QE03B - control group M = 4.3; test group M = 4.3; expert group M = 4.0). As expected, the game content was highly relevant for the experts, and 90.9% agreed or strongly agreed. However, 60% of participants in the control group and 73.9% of participants of the test group indicated that the game content is relevant with their interests as well (QE04A - control group M = 3.8; test group M = 3.9; expert group M = 4.1). For almost all study participants (96.6%) it was clear that the game contents are related to the human microbiome (QE04B - control group M = 4.6; test group M = 4.9; expert group M = 4.4), 83.1% agree that “Tiny Biome Tales” is an adequate teaching method for this subject (QE04C - control group M = 4.2; test group M = 4.3; expert group M = 4.3) and 72.3% also prefer learning with this game above other learning methods (QE04D - control group M = 4.0; test group M = 4.3; expert group M = 3.7). 72.3% of all individuals recruited for this pilot study had a satisfying feeling of accomplishment upon completing tasks in the game (QE05A - control group M = 4.2; test group M = 3.9; expert group M = 3.7) and 86.2% about the things they learned from the game as well (QE05C - control group M = 4.3; test group M = 4.3; expert group M = 4.0). Interestingly, only 50.8% of all study participants agreed with the statement “It is due to my personal effort that I managed to advance in the game” (QE05B - control group M = 3.7; test group M = 3.6; expert group M = 3.4). In the end, however, 89.2% of all players would recommend “Tiny Biome Tales” to people willing to learn about the human microbiome (QE05D - control group M = 4.4; test group M = 4.6; expert group M = 4.3).

**Fig. 4:**
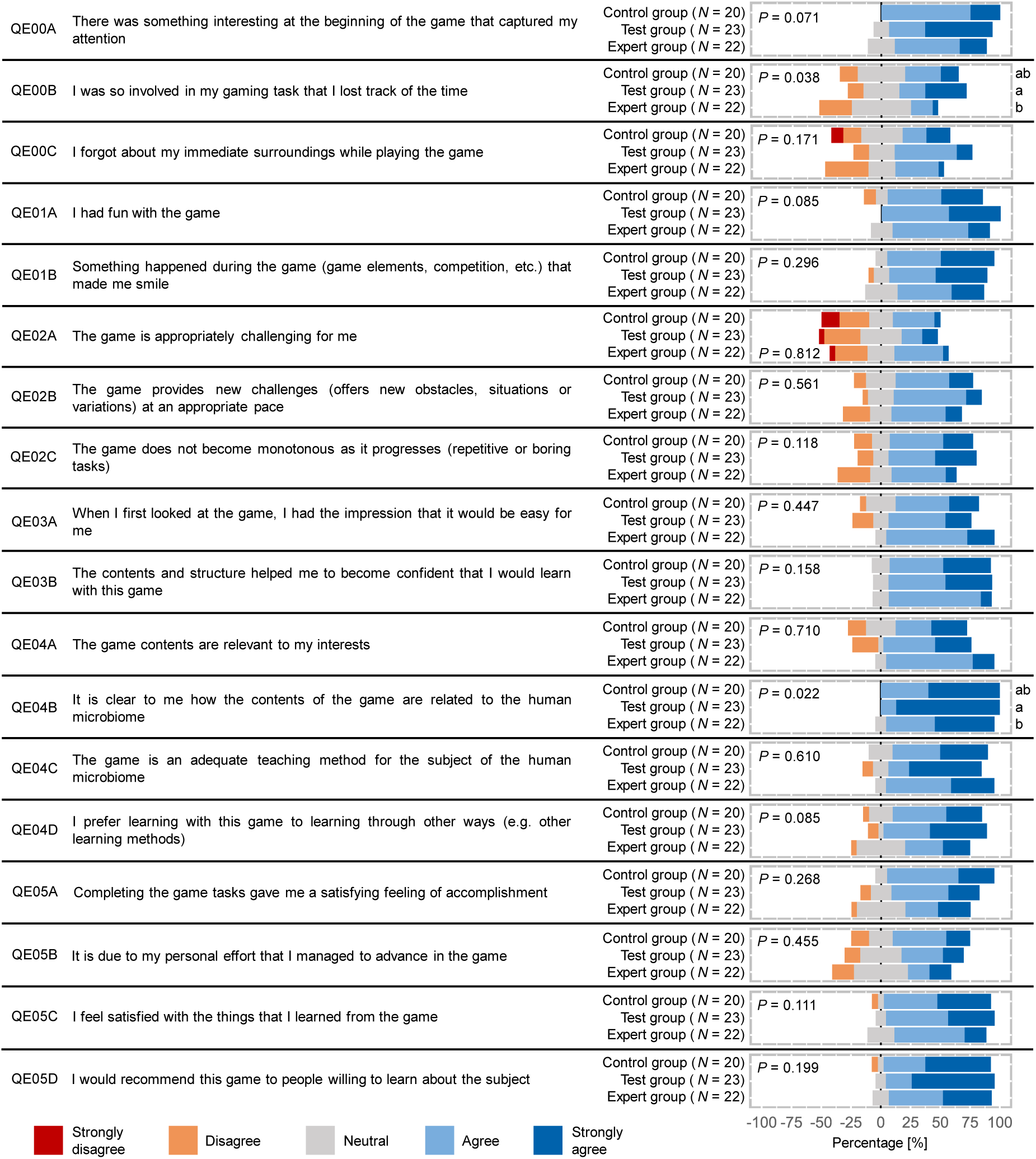
Player experience rating on a Likert scale from 1 (strongly disagree) to 5 (strongly agree). Ratings of the control group (*N* = 20), the test group (*N* = 23) and the expert group (*N* = 22) are each displayed in percent. Kruskal-Wallis test was performed to assess *P*-values. The letter code indicates significant differences (*P.adj.* 0.05 between groups, determined by Dunn‘s post hoc test with Benjamini-Hochberg correction of *P*-values.

### “Tiny Biome Tales contributed to my learning about the subject”

In addition to the player experience rating, the MEEGA+ model ^38^ provides for an assessment of usability including game aesthetics, learnability, operability and accessibility, and perceived learning (Fig. 5). Of all study participants, 72.3% agreed that the game design is visually attractive (QE06A - control group M = 3.8; test group M = 4.2; expert group M = 3.5) and 80% that the text font and colors are well blended and consistent (QE06B - control group M = 4.1; test group M = 4.5; expert group M = 3.8). A majority of players (75.4%) rated that they did not need to learn additional information before playing the game, suggesting that the game has intuitive gameplay (QE06C - control group M = 2.0; test group M = 1.8; expert group M = 2.1). Furthermore, 92.3%-98.5% of the study participants agreed that the game was easy to play, the game rules are clear and most people will find it easy to learn how to play “Tiny Biome Tales” (Q 06D- QE06G - control group M = 4.7-4.8; test group M = 4.6-4.9; expert group M = 4.1-4.3). The ratings by the expert group, however, were significantly lower in comparison to the control and test group, but still good with a mean of 4.1-4.3 (*P.adj.* < 0.05). Additionally, the expert group rated the readability of the font size and style (QE06I) significantly lower (*P.adj.* < 0.05) but still satisfactory, with a mean score of 4.1, in comparison to the control group (M = 4.6) and the test group (M = 4.7). 66.2% of all study participants agreed that the colors used in the game are meaningful (QE06J - control group M = 3.8; test group M =4.2; expert group M = 3.7). The game settings offer players the option to choose from three distinct color schemes: “green”, “red”, and “dark”. These selections alter various visual elements such as outline colors for selectable options, background colors for hover text, and basic UI components like buttons. Opting for the “dark” scheme additionally changes the text background color to black and the font color to white. Despite of these options for customization, only 44.6% of the players agreed that they could change the appearance according to their preference (QE06K - control group M = 3.6; test group M = 3.7; expert group M = 3.0). 5% of the control group and even 00% of the test group agreed that “Tiny Biome Tales” contributed to their learning about the human microbiome, and, although their rating was significantly lower (*P.adj.* < 0.05), also 81.2 of the test group agreed (QE07A - control group M = 4.4; test group M = 4.4; expert group M = 4.0). In the end, a majority of all study participants (81.5% agreed that “Tiny Biome Tales” allows for efficient learning about the human microbiome (QE07B - control group M = 4.1; test group M = 4.3; expert group M = 3.9).

**Fig. 5:**
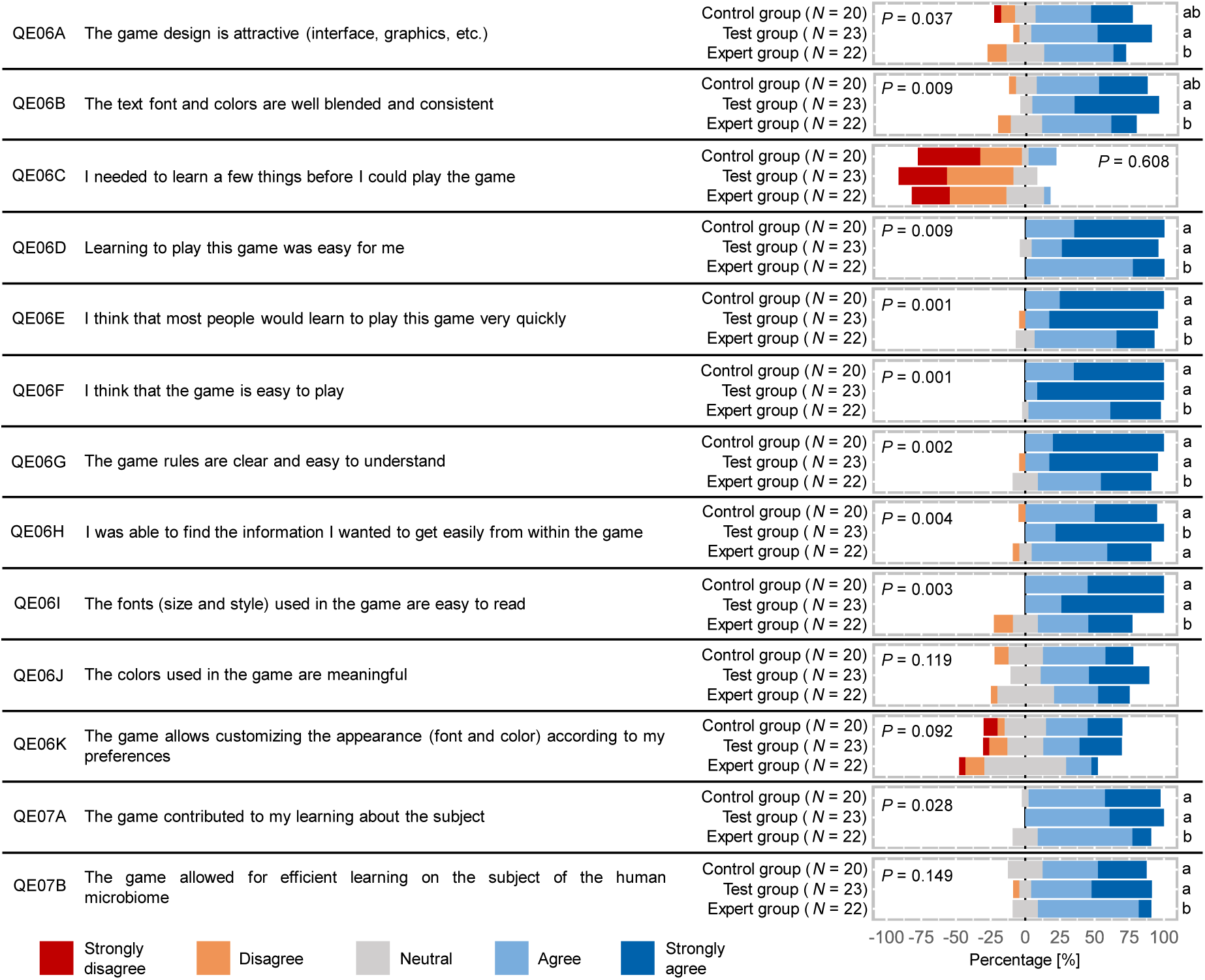
Usability and perceived learning rating on a Likert scale from 1 (strongly disagree) to 5 (strongly agree). Ratings of the control group (*N* = 20), the test group (*N* = 23) and the expert group (*N* = 22) are each displayed in percent. Kruskal-Wallis test was performed to assess *P*-values. The letter code indicates significant differences (*P.adj.* 0.05 between groups, determined by Dunn‘s post hoc test with Benjamini-Hochberg correction of *P*-values.

### “How small choices make a difference in life!”

Following the evaluation of player experience, usability, and perceived learning, all participants in the pilot study were given the opportunity to provide free text feedback in response to three questions (“Please list some good things about the game (if any)”, “Please give suggestions to improve the game (if any)”, “Any further comments?”; Supplementary Table 1-3). The feedback for each question was categorized into various themes, wherein each mention of a particular theme was counted to create a word cloud (Fig. 6).

**Fig. 6:**
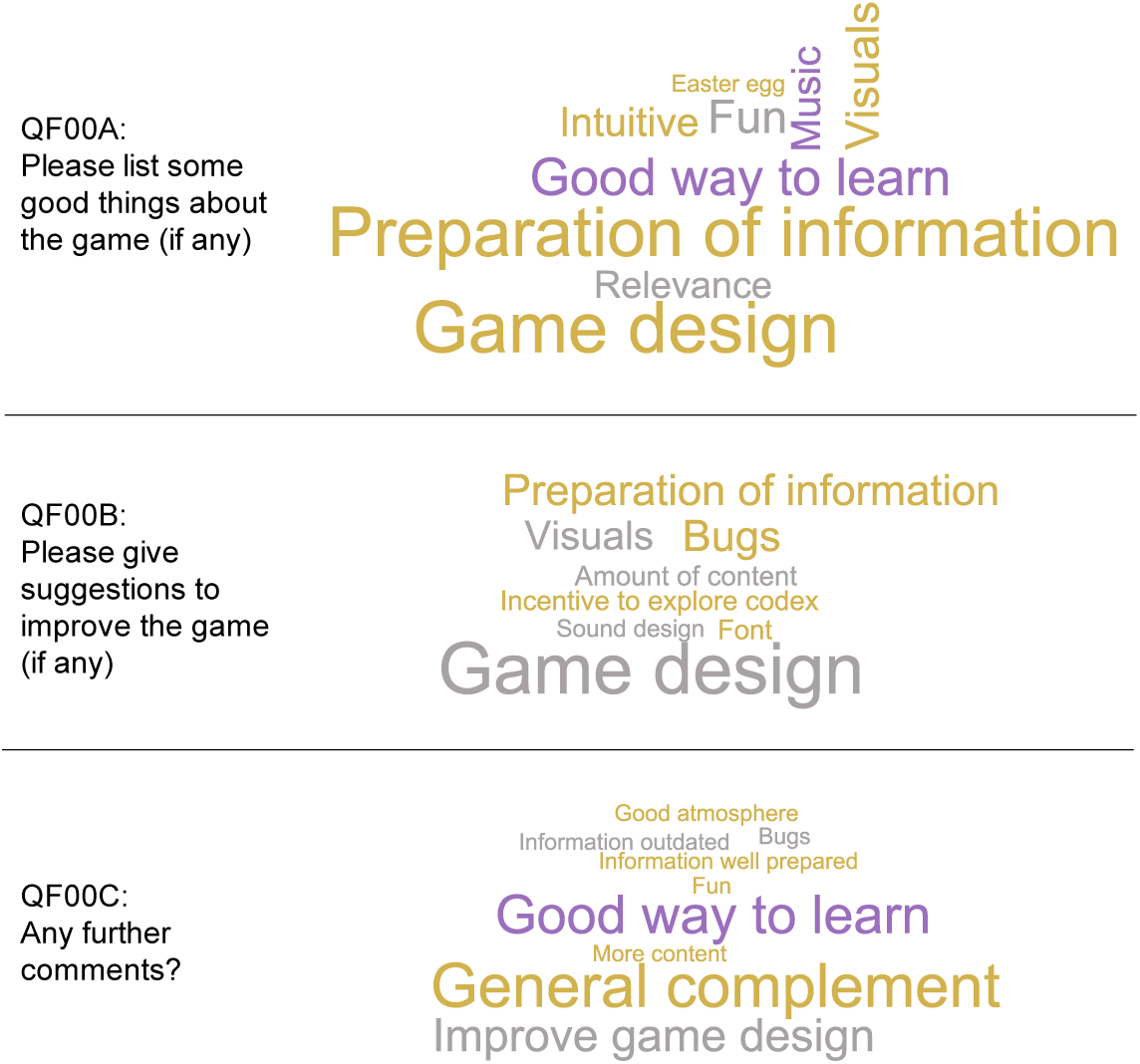
Word clouds displaying the frequency of generalized themes in the free text feedback. The colors indicate which group mentioned a particular theme the most: control group in gold, test group in purple and expert group in grey.

First, participants in the study were asked to list positive aspects of the game. Out of 65 participants, 45 used this opportunity. 42.2% of the 45 comments contained the theme “Game design” and 40% the theme “Preparation of information” praising the overall structure of the game and the clear presentation of information. Specifically, they appreciated the ability to explore topics of interest in depth and the feature of a diary where in-game decisions and their outcomes were repeated in more detail (Fig. 6 QF00A). Furthermore, frequently mentioned was that the game is “ un” (22.2% and playing it is a “Good way to learn” (26.7% about the human microbiome as well. Additionally, the quality of the “Music” (17.8%) and “ isuals” (22.2% were complemented, and players mentioned the “Relevance” (15.6% of the information: “How small choices make a difference in life!”

In response to the second open-text feedback question regarding potential improvements for the game, a total of 42 responses were received. It is noteworthy that the themes that were praised by many also are the ones that got the most criticism, with “Game design” (40.5%) emerging as the number one theme for improvement (Fig. 6 QF00B). Participants noted a lack of clarity regarding the required number of items to select in order to progress to the subsequent stage. Some participants expressed having to make choices that were not beneficial for the human microbiome due to a lack of alternative options. Furthermore, the text was found as overly complex and scientific (19%), particularly for individuals without prior knowledge of the human microbiome. Additionally, there were critiques regarding the “ isuals” (6.7% and several participants reported “Bugs” (19%), especially with disappearing text while playing in fullscreen mode.

Finally, 23 study participants used the opportunity to provide further comments. While some participants suggested to “Improve game design” (17. %), the majority of comments contained “General complements” (34.8%) and many participants commented here again that “Tiny Biome Tales” is a “Good way to learn” (26.1%) about the human microbiome (Fig. 6 QF00C).

## Discussion

The educational video game “Tiny Biome Tales” was developed with the primary intention to educate students and the public about the human microbiome, and to visualize the paradigm shift of microorganisms and disease towards microbiomes and health. To our knowledge, this is the first video game about the influence of lifestyle choices on the human microbiome and it’s relation to health. This pilot study demonstrated that playing “Tiny Biome Tales” significantly increased the knowledge about the human microbiome. The study participants agreed that the game allows for efficient learning and is an adequate teaching method on the subject. The content of the game was found relevant by most players. While playing, the study participants experienced fun and a satisfying feeling of accomplishment. The majority (98.5%) found the game intuitive to play and 89.2% of the study participants would recommend “Tiny Biome Tales” to people willing to learn about the human microbiome.

However, the primary objective for the player was to achieve the highest “Diversity Index” value, which may not align with the original development intention. For this particular objective, it appears that some players may have found it relatively straightforward to choose the correct options, as mentioned in the written feedback provided. The question arose whether there was a way to lose the game. In fact, there is not. This might explain why only 50.8% of the study participants agreed with the statement “It is due to my personal effort that I managed to advance in the game”. We assume that, even without prior knowledge of the human microbiome, most players will intuitively choose natural birth over cesarean section and breastfeeding over formula feeding in the game. Here, we also noted a bias in our questionnaire as the test group, although they played “Tiny Biome Tales” before answering the knowledge questionnaire, did not perform better in QK08 where we asked to which of the motheŕs body parts the microbiome of a cesarean born infant bears similarity. If not choosing “cesarean section” as a mode of delivery, players would never receive the information that the microbiome of a cesarean born infant resembles that of the mother’s skin surface ^42^. The control group performed slightly better compared to the test group at QK09 (“Is a decrease in short-chain fatty acid producing bacteria in the human gut associated with positive or negative health effects?”) as well. In this case, the information required was highly specific and may not have been relevant to the participant. To enhance the players’ learning motivation, it may be beneficial to incorporate a feature in the game that assesses the player’s knowledge about the microbiome, possibly in combination with a scoreboard, as wished by two study participants.

While the theme “Game design” received the most praise, it also garnered the most criticism in the open-text feedback. Some participants indicated that they occasionally felt compelled to select options that they believed to be detrimental to the microbiome, or wished for a return button to undo choices made accidentally, or did not get all the information they wanted because of bad game design. Also, three participants wished for more incentive to explore the “Codex”. In order to better align the game with its original intention of educating players about the human microbiome and also to address some of the major criticism, the primary focus of the game should be centered around acquiring knowledge instead of reaching the highest “Diversity Index”. However, our goal was to cater to a diverse audience, including those interested in delving deep into microbiome research as well as those seeking only entertaining gameplay with an overview on the microbiome, yet with key health information. This is indeed a balancing act that gives room for improving “Tiny Biome Tales”.

Furthermore, we understand that this game presents ideal lifestyle choices to players. We acknowledge that several options may not be equally available to all individuals all over the world due to various socio-economic factors. Also, choices for child delivery modes and newborn feeding are often not given due to medical or individual circumstances. Overall, we would like to clarify that we do not intend to promote an unrealistic and idealized view with perfect access to nutrition and healthcare. Instead, while developing the game we have focused on incorporating evidence from microbiome research to represent best practices for health in regard to the microbiome.

The open-text feedback theme “Preparation of information” was the second most frequent theme in also both positive and negative feedback. Some respondents appreciated the scientific writing style, while others found it overly complex and difficult to understand. In summary, “Tiny Biome Tales” evolved into a “gamified review” rather than an educational video game. Timmis et al. wrote that “microbiology literacy in society must become reality” ^27^. We are confident that once major flaws in game design and bugs are addressed, “Tiny Biome Tales” has the potential to serve as a platform for disseminating upcoming discoveries about the influence of daily habits and lifestyle choices on the human microbiome and potential health consequences in an engaging and accurate manner.

## Methods

### Study design

A pilot study was conducted for evaluating the quality and learning outcome of the educational video game “Tiny Biome Tales”. The study was performed fully online using the tool LimeSurvey (LimeSurvey GmbH, Hamburg, Germany) with no supervision from the creators. Participants were recruited by reaching out to colleagues from the field of microbiome research, posting to social media and via posters at Graz University of Technology. The time for playing the game was recorded and participants who had insufficient playtime to properly finish the game were excluded. In the end, we were able to recruit 65 individuals to play and evaluate “Tiny Biome Tales”. They were categorized into two main groups based on their prior knowledge about the human microbiome: the expert group (*N* = 22) and the non-expert group (*N* = 43). All study participants were instructed to complete demographic questions (Table 1) and a questionnaire based on the MEEGA+ model ^38, 39^ to rate their experience during playing on a Likert scale ranging from 1 (strongly disagree) to 5 (strongly agree) (Fig. 7). In order to evaluate the effectiveness of learning through playing the game, the non-experts were divided into two groups. The control group (*N* = 20) completed a knowledge questionnaire before playing the game, while the test group (*N* = 23) completed the same questionnaire after finishing the game.

### Game development

The game was developed using the Unity3D game engine (Unity Technologies, San Francisco, California, USA). WebGL (Khronos Group, Beaverton, Oregon, USA) was chosen as the deployment method. Low poly 3D assets from the POLYGON series (Synty Studios, Wellington, New Zealand) were purchased to build the scenes. The sound effects for the game were recorded using Audacity (Muse Group, Limassol, Cyprus) and a Shure SM58 microphone (Shure, Niles, Illinois, USA) in combination with a Scarlett 2i2 (Focusrite PLC, High Wycombe, England) interface. A Blue Yeti USB Microphone (Blue Microphones, Westlake Village, California, USA) was used as a secondary solution. The background music was crafted specifically for the game with FL Studio (Image-Line Software, Ghent, Belgium). All text content was created using information from the peer-reviewed articles that are referenced within the game.

### Statistics

All descriptive statistics and group comparisons via hypothesis testing were performed in R version 4.3.3 ^43^. Stacked bar plots and boxplots were generated using the ggplot2 package (version 3.5.0) ^44^, Likert plots with the likert package (version 2.0.0) ^45^, and word clouds using the wordcloud package (version 2.6) ^46^. To analyze if the test group outperformed the control group in the knowledge questionnaire, the wilcox.test and the fisher.test functions (R stats 4.3.3) were used to perform one-sided Mann–Whitney *U* tests and one-sided isher’s exact tests, respectively. The kruskal.test function (R stats 4.3.3) was used to execute Kruskal-Wallis rank sum tests to check if the participant groups rated the game differently and to check for differences in the demography between the groups in case of ordinal data, with post-hoc Dunn’s tests for pairwise comparison using the dunnTest function of the SA package (version 0.9.5) ^47^. To analyze differences between groups in the nominal demography data, isher’s exact tests with post-hoc pairwise comparison were utilized by executing the pairwiseNominalIndependence function of the rcompanion package (version 2.4.35) ^48^. In all pairwise comparisons, *P*-values were corrected with the Benjamini-Hochberg method.

## Acknowledgements

We express our gratitude to Tomislav Cernava for his insightful suggestions and discussions throughout the game development and creation of the MOOC. We appreciate Ahmed Abdellfattah for his expertise and contribution to the video production for the MOOC. Additionally, we would like to acknowledge the assistance provided by Stefan Janisch, Antonios Rouvelas, and the iMooX team in supporting the development of the MOOC. The developed work presented here was co-funded by the Federal Ministry of Education, Science and Research, Austria, as part of the 2019 call for proposals for digital and social transformation in higher education for the project “iMooX - die MOOC-Plattform als Service für alle österreichischen Hochschulen” (2021-2023, partner organisations: University of Vienna, Graz University of Technology).

