## Supplementary tables 1-3 for "“Tiny Biome Tales”: a gamified review about the influence of lifestyle choices on the human microbiome"

Supplementary table 1: **Answers to the question QF00A - Please list some good things about the game (if any).** The free text feedback of all three groups (control group - *N* = 20, test group - *N* = 23 and expert group - *N* = 22) was categorized into topics.

| | | Counts of complements on following topics: | | | | | | | | |
|---|---|---|---|---|---|---|---|---|---|---|
| **Participant ID** | **Free text feedback** | **Relevance** | **Intuitive** | **Preparation of information** | **Music** | **Visuals** | **Game design** | **Good way to learn** | **Easter egg** | **Fun** |
| | **Control group (*N* = 20)** | | | | | | | | | |
| 3 | "The choices were meaningful, because of their relevance in everyday life" | 1 | 0 | 0 | 0 | 0 | 0 | 0 | 0 | 0 |
| 4 | "i liked how easy it was to understand and play this game. the colors and sounds were also to my liking. i also liked how much information you got after every ,,level"" and that you got to read the information any time within the game." | 0 | 1 | 1 | 0 | 0 | 0 | 0 | 0 | 0 |
| 5 | [German] "klar verständlich, jugendfrei" | 0 | 0 | 1 | 0 | 0 | 0 | 0 | 0 | 0 |
| 6 | "Nice music and nice art style. I like that I can view my decisions in retrospective. The game gives a very chill vibe. 10/10." | 0 | 0 | 0 | 1 | 1 | 1 | 0 | 0 | 0 |
| 7 | "The game is easy to learn, which makes it simple to just give it a go. Also the information, the game is giving you, is great. Even if most information is just presented in textual form you get the feeling for the meaning of your actions for your microbiome." | 0 | 1 | 1 | 0 | 0 | 1 | 0 | 0 | 0 |
| 9 | "Really inuitive, easy handling, the + and - for the feedback are nice. Also the sum up at the end was helpful to repeat the impacts." | 0 | 1 | 0 | 0 | 0 | 1 | 0 | 0 | 0 |
| 10 | N/A | 0 | 0 | 0 | 0 | 0 | 0 | 0 | 0 | 0 |

| | | | | | | | | | | |
|---|---|---|---|---|---|---|---|---|---|---|
| 11 | "The graphics and music made it pleasant to spend time with the game, it did not feel like I was sitting down to learn something but the learning part came as a sort of reward for also performing the actions you thought were right - or wrong. I really liked the ""wiki"" section that allowed you to re read certain information and also made the sources easily accessible." | 0 | 0 | 0 | 1 | 1 | 1 | 1 | 0 | 0 |
| 12 | "Funny and amusing design, suitable font size and font, short explanations -> not too much words" | 0 | 0 | 1 | 0 | 1 | 0 | 0 | 0 | 0 |
| 15 | "easy introductions, very informative and well cited, great design" | 0 | 1 | 1 | 0 | 0 | 1 | 0 | 0 | 0 |
| 19 | "Pickle RIIIIIICK." | 0 | 0 | 0 | 0 | 0 | 0 | 0 | 1 | 0 |
| 20 | "Easter eggs and it was easy to understand " | 0 | 0 | 1 | 0 | 0 | 0 | 0 | 1 | 0 |
| 22 | "consistent + cohesive layout and design, nice interactions relatively cool sand castle great way of learning about microbiomes" | 0 | 0 | 0 | 0 | 1 | 1 | 1 | 0 | 0 |
| 24 | "Nice game design, witty choice of daily activities to involve to lean about the human microbiom" | 1 | 0 | 0 | 0 | 0 | 1 | 0 | 0 | 0 |
| 26 | "Good summary of information. Liked that you made choices yourself" | 0 | 0 | 1 | 0 | 0 | 1 | 0 | 0 | 0 |
| 27 | "simple user-interface, good storyline, actions and consequences are explained very well, the soundtrack is very nice and relaxing, the information interface with embedded sources is well done" | 0 | 1 | 1 | 1 | 0 | 1 | 0 | 0 | 0 |
| 28 | [German] "schönes Design, hat mir gut gefallen, habe einiges dazugelernt" | 0 | 0 | 0 | 0 | 1 | 0 | 1 | 0 | 0 |
| 29 | N/A | 0 | 0 | 0 | 0 | 0 | 0 | 0 | 0 | 0 |
| 30 | "issa funny game ,Sometimes the texts didnt show up :/" | 0 | 0 | 0 | 0 | 0 | 0 | 0 | 0 | 1 |
| 32 | N/A | 0 | 0 | 0 | 0 | 0 | 0 | 0 | 0 | 0 |
| **Test group (*N* = 23)** | | | | | | | | | | |

| | | | | | | | | | | |
|---|---|---|---|---|---|---|---|---|---|---|
| 4 | "It was very informative and made it interesting to learn new things about the human microbiome." | 0 | 0 | 1 | 0 | 0 | 0 | 1 | 0 | 0 |
| 5 | "Very coherent and fun to interact with. Directly reflects on the choices made by giving information about the actions backed with studies etc. Revealed interesting facts about bacteria in general and how it affects our microbiome." | 1 | 0 | 0 | 0 | 0 | 1 | 0 | 0 | 1 |
| 7 | "Instructive" | 0 | 0 | 0 | 0 | 0 | 0 | 1 | 0 | 0 |
| 8 | "very calming music track, not distracting or strenuous" | 0 | 0 | 0 | 1 | 0 | 0 | 0 | 0 | 0 |
| 10 | "Fun!" | 0 | 0 | 0 | 0 | 0 | 0 | 0 | 0 | 1 |
| 11 | "Based on actual papers and directly linkedt to them" | 0 | 0 | 0 | 0 | 0 | 1 | 0 | 0 | 0 |
| 14 | "Story, Sounds, Atmosphere" | 0 | 0 | 0 | 0 | 0 | 1 | 0 | 0 | 0 |
| 15 | "the colors, fonts and graphics make it comfortable to play, and i learned some things about pregnancy and childbirth i didn't know before" | 0 | 0 | 0 | 0 | 1 | 0 | 1 | 0 | 0 |
| 16 | N/A | 0 | 0 | 0 | 0 | 0 | 0 | 0 | 0 | 0 |
| 18 | N/A | 0 | 0 | 0 | 0 | 0 | 0 | 0 | 0 | 0 |
| 19 | N/A | 0 | 0 | 0 | 0 | 0 | 0 | 0 | 0 | 0 |
| 21 | "lovely details in the background ect., backgroundmusic is nice, very detailed information, i really like it!" | 0 | 0 | 1 | 1 | 1 | 0 | 0 | 0 | 0 |
| 22 | N/A | 0 | 0 | 0 | 0 | 0 | 0 | 0 | 0 | 0 |
| 23 | "fucking beautiful, good length, replay value" | 0 | 0 | 0 | 0 | 1 | 1 | 0 | 0 | 0 |
| 24 | "Very easy to understand. Nice explanation of the effects that the actions you take have. Music and sounds are relaxing and not disturbing. Smooth to play." | 0 | 1 | 1 | 1 | 0 | 1 | 0 | 0 | 0 |
| 25 | N/A | 0 | 0 | 0 | 0 | 0 | 0 | 0 | 0 | 0 |
| 26 | N/A | 0 | 0 | 0 | 0 | 0 | 0 | 0 | 0 | 0 |
| 27 | N/A | 0 | 0 | 0 | 0 | 0 | 0 | 0 | 0 | 0 |
| 28 | "Easy to play, informative, good alternative to social media" | 0 | 1 | 0 | 0 | 0 | 0 | 1 | 0 | 0 |
| 31 | "Music is chill" | 0 | 0 | 0 | 1 | 0 | 0 | 0 | 0 | 0 |
| 32 | "easy to understand (basic language), fun graphics, makes studying a little bit more fun" | 0 | 0 | 1 | 0 | 1 | 0 | 1 | 0 | 0 |

| | | | | | | | | | | |
|---|---|---|---|---|---|---|---|---|---|---|
| 37 | "Great game to learn (more) about the human microbiome in an easier way. Good explanations of the respective situations in realtion to the human microbiome. Had a lot of fun playing through different scenarios and learning new things every time." | 0 | 0 | 1 | 0 | 0 | 0 | 1 | 0 | 1 |
| 39 | "It is always fun to play a game and to learn new things! It was a fun game and I learnd a lot!" | 0 | 0 | 0 | 0 | 0 | 0 | 1 | 0 | 1 |
| | **Expert group (*N* = 22)** | | | | | | | | | |
| 5 | "I already like the game. Very engaging, personally for me it was nice to be a pregnant woman for a change (a brief detour from a man's life). Good things, in my opinion, are as follows: 1. Teaching with real-life scenarios (for Eg. what would you do at a party post-pregnancy (drink/dance), what would you eat at a picnic, etc); 2. The recap or the effect of my decisions throughout the day/scenes was a nice feature. It accomplishes 2 things a. You get the direct consequence of your decisions (like an increase in diversity post-exercise, Bacteroidetes-Firmicutes ratio, etc) immediately after every scene. It helps to make better choices as the game progresses. b. You reinforce the learnings associated with those actions; 3. The music and graphics are just enough to engage and not distract too much from the scenes" | 0 | 0 | 0 | 1 | 1 | 1 | 1 | 0 | 0 |
| 7 | [German] "Sehr geringe Einstiegshürde, sehr umfangreiche und detaillierte Erklärungen, liebe zum Detail" | 0 | 1 | 1 | 0 | 0 | 0 | 0 | 0 | 0 |
| 8 | "fantastic idea, very creative" | 0 | 0 | 0 | 0 | 0 | 1 | 0 | 0 | 0 |
| 9 | N/A | 0 | 0 | 0 | 0 | 0 | 0 | 0 | 0 | 0 |
| 11 | "awesome work, i like that some relevant information is repeated before the next chapter, sometimes I missed the plus/minus, so it was nice to have it summarized again" | 0 | 0 | 0 | 0 | 0 | 1 | 0 | 0 | 0 |

| | | | | | | | | | | |
|---|---|---|---|---|---|---|---|---|---|---|
| 12 | "immediate start ("I got pregnant" - hilarious!), Very well investigated, Scientific references available if someone is interested, Life history of a human" | 1 | 0 | 1 | 0 | 0 | 0 | 0 | 0 | 1 |
| 13 | N/A | 0 | 0 | 0 | 0 | 0 | 0 | 0 | 0 | 0 |
| 19 | N/A | 0 | 0 | 0 | 0 | 0 | 0 | 0 | 0 | 0 |
| 20 | "It is a good collection of references and summaries of important events. I liked the disclosure about the diversity index from the beginning but one plays the entire game without knowing what would be a good diversity index throughout the different life stages." | 0 | 0 | 1 | 0 | 0 | 0 | 0 | 0 | 0 |
| 21 | "It was fun playing the game and I would recommend it to anyone that would like to learn about the microbiome. The information was not too less and not too much and easy to understand." | 0 | 0 | 1 | 0 | 0 | 0 | 1 | 0 | 1 |
| 22 | N/A | 0 | 0 | 0 | 0 | 0 | 0 | 0 | 0 | 0 |
| 23 | N/A | 0 | 0 | 0 | 0 | 0 | 0 | 0 | 0 | 0 |
| 24 | N/A | 0 | 0 | 0 | 0 | 0 | 0 | 0 | 0 | 0 |
| 27 | "The information provided are very educative & interesting. It's really fun to have different experience and get different information when I made different choice in the game" | 0 | 0 | 1 | 0 | 0 | 1 | 0 | 0 | 1 |
| 28 | N/A | 0 | 0 | 0 | 0 | 0 | 0 | 0 | 0 | 0 |
| 30 | N/A | 0 | 0 | 0 | 0 | 0 | 0 | 0 | 0 | 0 |
| 31 | N/A | 0 | 0 | 0 | 0 | 0 | 0 | 0 | 0 | 0 |
| 34 | "I guess this is a very good initial effort to introduce the significance of microbiome diversity knowledge in our everyday choices. Sometimes, its easy to forget as a researcher that this topic should be realised in everyday life." | 1 | 0 | 0 | 0 | 0 | 0 | 0 | 0 | 0 |
| 40 | "egaging, very fun to progress throughoud life periods of an individual person and the interactions that someone encounters" | 1 | 0 | 0 | 0 | 0 | 0 | 0 | 0 | 1 |

| | | | | | | | | | | |
|---|---|---|---|---|---|---|---|---|---|---|
| 41 | "The game did captivate my interest and it was entertaining. I also learned some facts I didn't know, and made me want to look more into some information." | 0 | 0 | 0 | 0 | 0 | 1 | 0 | 0 | 1 |
| 43 | N/A | 0 | 0 | 0 | 0 | 0 | 0 | 0 | 0 | 0 |
| 45 | "How small choices make a difference in life! " | 1 | 0 | 0 | 0 | 0 | 0 | 0 | 0 | 0 |

N/A: no answer given by the study participant

Supplementary table 2: **Answers to the question QF00B - Please give suggestions to improve the game (if any).** The free text feedback of all three groups (control group - *N* = 20, test group - *N* = 23 and expert group - *N* = 22) was categorized into topics.

| | | **Counts of mentions of the following topics that need improvement** | | | | | | | |
|---|---|---|---|---|---|---|---|---|---|
| **Participant ID** | **Free text feedback** | **Amount of content** | **Font** | **Visuals** | **Preparation of information** | **Game design** | **Incentive to explore codex** | **Bugs** | **Sound design** |
| | **Control group (*N* = 20)** | | | | | | | | |
| 3 | "add more scenes" | 1 | 0 | 0 | 0 | 0 | 0 | 0 | 0 |
| 4 | "the font was a little to small." | 0 | 1 | 0 | 0 | 0 | 0 | 0 | 0 |
| 5 | [German] "Grafik, kürzere Erklärungen (mehr Stichwörter), selbst entscheiden wie viele Sachen man auswählt" | 0 | 0 | 1 | 1 | 1 | 0 | 0 | 0 |
| 6 | "Maybe use another font. The information tab was relatively unused by me during the game but I like the idea. It would be very interesting if one could read all informations after the game and not only the unlocked ones." | 0 | 1 | 0 | 0 | 0 | 1 | 0 | 0 |
| 7 | "The game can be a bit stale with all information just being text. So the first thing I would try to implement, would be more diverse ways of presenting all the information. Aside from that the unlocks are something I did not use at all because I just did not notice them. This could be changed as well." | 0 | 0 | 0 | 1 | 0 | 1 | 0 | 0 |
| 9 | "A possibility to return to a previous point, when accidentally clicking wrong. A bit of overloading of information " | 0 | 0 | 0 | 0 | 1 | 0 | 0 | 0 |

| | | | | | | | | | |
|---|---|---|---|---|---|---|---|---|---|
| 10 | "During the end (partyscene, kitchen) the game began to lag and my Laptop had issues handling the game. Other then that the game was really fun to play" | 0 | 0 | 0 | 0 | 0 | 0 | 1 | 0 |
| 11 | "For some decisions I wish there was more of a choice or different actions you can take." | 0 | 0 | 0 | 0 | 1 | 0 | 0 | 0 |
| 12 | "Maybe include more hints on how to get to the next step --> Sometimes it wasn't clear what you had to click on to get to the next step" | 0 | 0 | 0 | 0 | 1 | 0 | 0 | 0 |
| 15 | "didn't know how often I have to click when at the table with food (for example) to continue with the game... I thought I only have to choose one item" | 0 | 0 | 0 | 0 | 1 | 0 | 0 | 0 |
| 19 | N/A | 0 | 0 | 0 | 0 | 0 | 0 | 0 | 0 |
| 20 | N/A | 0 | 0 | 0 | 0 | 0 | 0 | 0 | 0 |
| 22 | "it's interesting to learn about the microbiome and its specific vocab, but the overload of technical terms can be overwhelming for newbies i think the texts would be easier to read if a different quotation guideline was used" | 0 | 0 | 0 | 1 | 0 | 0 | 0 | 0 |
| 24 | "very educational, but maybe sometimes a little hard to understand for a non-academical and/or non-native/non fluent English speaker" | 0 | 0 | 0 | 1 | 0 | 0 | 0 | 0 |
| 26 | "Add a layer of complexity. Very linear progression" | 0 | 0 | 0 | 0 | 1 | 0 | 0 | 0 |
| 27 | "has quite a few bugs (sometimes font does not appear, when exiting full screen mode with esc and reentering full screen mode, a black screen appears always, sometimes the game would not load, I had to reopen it in another window) It would be a great improvement if you could exit certain interfaces (like the ones giving information, and the progress book) with esc" | 0 | 0 | 0 | 0 | 0 | 0 | 1 | 0 |

| | | | | | | | | | |
|---|---|---|---|---|---|---|---|---|---|
| 28 | [German] "Wenn man den Vollbildmodus einschaltet, funktionieren die Anweisungen nicht mehr und das Spiel muss neu gestartet werden, um die Anweisungen wieder zu sehen." | 0 | 0 | 0 | 0 | 0 | 0 | 1 | 0 |
| 29 | N/A | 0 | 0 | 0 | 0 | 0 | 0 | 0 | 0 |
| 30 | N/A | 0 | 0 | 0 | 0 | 0 | 0 | 0 | 0 |
| 32 | N/A | 0 | 0 | 0 | 0 | 0 | 0 | 0 | 0 |
| | **Test group (*N* = 23)** | | | | | | | | |
| 4 | N/A | 0 | 0 | 0 | 0 | 0 | 0 | 0 | 0 |
| 5 | "Click sound when choosing the Items" | 0 | 0 | 0 | 0 | 0 | 0 | 0 | 1 |
| 7 | "Personally I think these texts are a little bit too academic. That makes it optmial for students and university absolvents but harder for the ""normal"" people." | 0 | 0 | 0 | 1 | 0 | 0 | 0 | 0 |
| 8 | "more incentives to read up in the in-game wiki" | 0 | 0 | 0 | 0 | 0 | 1 | 0 | 0 |
| 10 | "Sometimes you had to choose something with negative effect to be able to leave the situation." | 0 | 0 | 0 | 0 | 1 | 0 | 0 | 0 |
| 11 | If you don't know the basics (like what is a microbiom, what are microorganism, how can they be classified how do they work...) then I guess it is a bit hard to follow the explanations | 0 | 0 | 0 | 1 | 0 | 0 | 0 | 0 |
| 14 | N/A | 0 | 0 | 0 | 0 | 0 | 0 | 0 | 0 |
| 15 | N/A | 0 | 0 | 0 | 0 | 0 | 0 | 0 | 0 |
| 16 | N/A | 0 | 0 | 0 | 0 | 0 | 0 | 0 | 0 |
| 18 | N/A | 0 | 0 | 0 | 0 | 0 | 0 | 0 | 0 |
| 19 | N/A | 0 | 0 | 0 | 0 | 0 | 0 | 0 | 0 |
| 21 | N/A | 0 | 0 | 0 | 0 | 0 | 0 | 0 | 0 |
| 22 | N/A | 0 | 0 | 0 | 0 | 0 | 0 | 0 | 0 |
| 23 | "the codex feels a bit overwhelming at first sight" | 0 | 0 | 0 | 1 | 0 | 0 | 0 | 0 |
| 24 | "The actions you should take for positive affects are too recognizable. (Maybe not the goal of the game to be hard but a little bit more challenging would be nice for me)" | 0 | 0 | 0 | 0 | 1 | 0 | 0 | 0 |

| | | | | | | | | | |
|---|---|---|---|---|---|---|---|---|---|
| 25 | "There is a lot of text to read. Sometimes it would be great if it was broken down a bit more" | 0 | 0 | 0 | 1 | 0 | 0 | 0 | 0 |
| 26 | N/A | 0 | 0 | 0 | 0 | 0 | 0 | 0 | 0 |
| 27 | "The overall result at the end of the game was a little ambigous. ""[...]Most of your decisions had a very positive effect on your health. You made very few decision which had a positive effect of your health[...]"". My lifestyle was rather unhealthy during that playthrough." | 0 | 0 | 0 | 0 | 0 | 0 | 1 | 0 |
| 28 | N/A | 0 | 0 | 0 | 0 | 0 | 0 | 0 | 0 |
| 31 | "Scoreboard." | 0 | 0 | 0 | 0 | 1 | 0 | 0 | 0 |
| 32 | "some of the explanation brackets have no words written in them (for example during tutorial)" | 0 | 0 | 0 | 0 | 0 | 0 | 1 | 0 |
| 37 | "In full screen mode, some texts and overlays could not be displayed. If the game was not played in full sreen mode, all text elements were visible." | 0 | 0 | 0 | 0 | 0 | 0 | 1 | 0 |
| 39 | N/A | 0 | 0 | 0 | 0 | 0 | 0 | 0 | 0 |
| **Expert group (*N* = 22)** | | | | | | | | | |
| 5 | "Maybe add more stories/scenes for kids to play (starting with pregnancy for kids may be a little boring/awkward). In future updates, for kids maybe a different storyline can be added. Like, 1. During a school party, what would you eat/drink? 2. In a free period/class what activities can be done (gardening/playing games/ sitting or doing nothing and eating sugary candy) 3. When you are hungry and you go to the supermarket with your parents/alone what would you buy? etc. Maybe this also applies to older adults." | 1 | 0 | 0 | 0 | 0 | 0 | 0 | 0 |
| 7 | "Für mich war teilweise nicht klar wie viele Optionen zum Vorankommen ausgewählt werden müssen." | 0 | 0 | 0 | 0 | 1 | 0 | 0 | 0 |

| | | | | | | | | | |
|---|---|---|---|---|---|---|---|---|---|
| 8 | "if you are interested in gaming, the content and appearance of the game seems a bit outdated (graphics, simple point and click), simple audio, no videos, no animations, very static" | 0 | 0 | 1 | 0 | 1 | 0 | 0 | 1 |
| 9 | N/A | 0 | 0 | 0 | 0 | 0 | 0 | 0 | 0 |
| 11 | "the plus and minus could be a bit more prominent" | 0 | 0 | 1 | 0 | 0 | 0 | 0 | 0 |
| 12 | "Highscore for players, expand, Easter eggs/""sidequests"" (maybe finding specific items?), Graphics of the humans (especially friend at picnic), Adding some random events that the player can not deterministically influence the outcome" | 0 | 0 | 1 | 0 | 1 | 0 | 0 | 0 |
| 13 | N/A | 0 | 0 | 0 | 0 | 0 | 0 | 0 | 0 |
| 19 | N/A | 0 | 0 | 0 | 0 | 0 | 0 | 0 | 0 |
| 20 | "I know the game intends to be informative but it puts a weird pressure on every daily action. I guess that maybe if it wasn't called a game but more like an interactive exploration of the human microbiome in our daily lives I would be less frustrating to have to pick something I didn't want to, eg: I actually didn't want to pick pizza at the beginning but at some point, it was either that or cigarettes." | 0 | 0 | 0 | 0 | 1 | 0 | 0 | 0 |
| 21 | "It was not clear to me that I was aging during the game. It was said that I am 2 years old and then I went to a party and had to clean my apartment. It would be nice if the aging process could be stated. Sometimes I did not get information about other things I would like to know, e.g. "my friend" was eating the veg. wrap at the picnic and I had no chance to get information about the influence on the microbiome." | 0 | 0 | 0 | 0 | 1 | 0 | 0 | 0 |
| 22 | N/A | 0 | 0 | 0 | 0 | 0 | 0 | 0 | 0 |
| 23 | N/A | 0 | 0 | 0 | 0 | 0 | 0 | 0 | 0 |
| 24 | N/A | 0 | 0 | 0 | 0 | 0 | 0 | 0 | 0 |

| | | | | | | | | | |
|---|---|---|---|---|---|---|---|---|---|
| 27 | "Please improve the graphic design & the color" | 0 | 1 | 1 | 0 | 0 | 0 | 0 | 0 |
| 28 | "when choosing the drinks to bring for the picnic, you couldn't choose something healthy / a pregnant woman would take - I assume no pregnant woman would bring coffee or beer. At the picnic or the barbecue you had to choose something clearly unhealthy which was not good for your microbiome as you couldn't continue until you chose something else" | 0 | 0 | 0 | 0 | 1 | 0 | 0 | 0 |
| 30 | "it's interesting with good information however the texts are sometimes too long ,make the information more concise" | 0 | 0 | 0 | 0 | 0 | 0 | 0 | 0 |
| 31 | N/A | 0 | 0 | 0 | 0 | 0 | 0 | 0 | 0 |
| 34 | "I would like more animation please rather than a text with journal citation, a documentary video that illustrate the points would be a super good addition to the game" | 0 | 0 | 1 | 0 | 0 | 0 | 0 | 0 |
| 40 | "to have the title of a life period in a setting written, or at least introduced - like the stage: pregnancy and birth. Maybe it is, but none of the popups in the game has showed on my screen, only after the completion of each level i say the book and the summary of all the decisions. firts part with choices on the cesarean section vs natural birth and the mode of feeding the baby are actually often not a choice of the mother and cannot be treated as such. The C section is often pushed by clinitians and it is done in the case of emergencies.. so it is not a choice that one can deliberately make. the same is with breastfeeding. So, i would advise to put some cautionary, gentle comments in that text, as you did with antibiotics. As of now, it sounds harsh. It is not the same thing as with choosing what to eat, or as to brush your teeth. The birth and breastfeeding are often connected with the necessity and medical decisions" | 0 | 0 | 0 | 0 | 1 | 0 | 1 | 0 |

| | | | | | | | | | |
|---|---|---|---|---|---|---|---|---|---|
| 41 | "After being a 2 years old, I didn't see that i turned into an adult again, and suddenly I was in a club, so that was a bit confusing. Also, when I turned the game full screen I couldn't see the text (the squares were blank) and I had to refresh the page and start again. On the part where the information about the bacteria is, it would be good to have some basic facts listed at the beginning so you can understand quickly without having to read the whole text (for example: relevance (high to low), if its beneficial or not, stage of life were its relevant, diseases related, etc.) and if you are interested to know more details then you can also read the text." | 0 | 0 | 0 | 0 | 1 | 0 | 1 | 0 |
| 43 | "The graphic could be better" | 0 | 0 | 1 | 0 | 0 | 0 | 0 | 0 |
| 45 | "More characters could be included in more diverse situations. " | 1 | 0 | 0 | 0 | 0 | 0 | 0 | 0 |

N/A: no answer given by the study participant

Supplementary table 3: **Answers to the question QF00C - Any further comments?** The free text feedback of all three groups (control group - *N* = 20, test group - *N* = 23 and expert group - *N* = 22) was categorized into topics.

| | | Counting mentions of the following topics | | | | | | | | |
|---|---|---|---|---|---|---|---|---|---|---|
| **Participant ID** | **Free text feedback** | **More content** | **Good atmosphere** | **Improve game design** | **Good way to learn** | **Information well prepared** | **General complement** | **Fun** | **Bugs** | **Information outdated** |
| | | **Control group (*N* = 20)** | | | | | | | | |
| 3 | If you extend the pregnancy part, this game could help couples to make the right choices during pregnancy | 1 | 0 | 0 | 0 | 0 | 0 | 0 | 0 | 0 |
| 4 | N/A | 0 | 0 | 0 | 0 | 0 | 0 | 0 | 0 | 0 |
| 5 | N/A | 0 | 0 | 0 | 0 | 0 | 0 | 0 | 0 | 0 |

| | | | | | | | | | | |
|---|---|---|---|---|---|---|---|---|---|---|
| 6 | The game scenarios were pretty well made. Many situations feel very real. For example at the beginning of the game in the living room where you can see the garden through the window and busses driving at the street. | 0 | 1 | 0 | 0 | 0 | 0 | 0 | 0 | 0 |
| 7 | N/A | 0 | 0 | 0 | 0 | 0 | 0 | 0 | 0 | 0 |
| 9 | "Some options felt mandatory in order to progress in the game" | 0 | 0 | 1 | 0 | 0 | 0 | 0 | 0 | 0 |
| 10 | N/A | 0 | 0 | 0 | 0 | 0 | 0 | 0 | 0 | 0 |
| 11 | "I think that games like these are a very good way of teaching about certain subjects in a playful way, especially for kids, and could also be used for other important topics to make learning feel less like a chore." | 0 | 0 | 0 | 1 | 0 | 0 | 0 | 0 | 0 |
| 12 | "Very well designed and good work - must have been a lot of preparation work" | 0 | 0 | 0 | 0 | 1 | 0 | 0 | 0 | 0 |
| 15 | "great length of the game, doesn't get boring!" | 0 | 0 | 0 | 0 | 0 | 1 | 0 | 0 | 0 |
| 19 | N/A | 0 | 0 | 0 | 0 | 0 | 0 | 0 | 0 | 0 |
| 20 | N/A | 0 | 0 | 0 | 0 | 0 | 0 | 0 | 0 | 0 |
| 22 | "fun game " | 0 | 0 | 0 | 0 | 0 | 0 | 1 | 0 | 0 |
| 24 | "nice game, great topic for a thesis" | 0 | 0 | 0 | 0 | 0 | 1 | 0 | 0 | 0 |
| 26 | N/A | 0 | 0 | 0 | 0 | 0 | 0 | 0 | 0 | 0 |
| 27 | "Good luck on your project, I´m sure it will turn out great :)" | 0 | 0 | 0 | 0 | 0 | 1 | 0 | 0 | 0 |
| 28 | N/A | 0 | 0 | 0 | 0 | 0 | 0 | 0 | 0 | 0 |
| 29 | N/A | 0 | 0 | 0 | 0 | 0 | 0 | 0 | 0 | 0 |
| 30 | N/A | 0 | 0 | 0 | 0 | 0 | 0 | 0 | 0 | 0 |
| 32 | N/A | 0 | 0 | 0 | 0 | 0 | 0 | 0 | 0 | 0 |
| | | **Test group (*N* = 23)** | | | | | | | | |
| 4 | N/A | 0 | 0 | 0 | 0 | 0 | 0 | 0 | 0 | 0 |
| 5 | N/A | 0 | 0 | 0 | 0 | 0 | 0 | 0 | 0 | 0 |
| 7 | N/A | 0 | 0 | 0 | 0 | 0 | 0 | 0 | 0 | 0 |
| 8 | N/A | 0 | 0 | 0 | 0 | 0 | 0 | 0 | 0 | 0 |

| | | | | | | | | | | |
|---|---|---|---|---|---|---|---|---|---|---|
| 10 | N/A | 0 | 0 | 0 | 0 | 0 | 0 | 0 | 0 | 0 |
| 11 | N/A | 0 | 0 | 0 | 0 | 0 | 0 | 0 | 0 | 0 |
| 14 | "Good Theme to bring up to the people" | 0 | 0 | 0 | 1 | 0 | 0 | 0 | 0 | 0 |
| 15 | N/A | 0 | 0 | 0 | 0 | 0 | 0 | 0 | 0 | 0 |
| 16 | N/A | 0 | 0 | 0 | 0 | 0 | 0 | 0 | 0 | 0 |
| 18 | N/A | 0 | 0 | 0 | 0 | 0 | 0 | 0 | 0 | 0 |
| 19 | N/A | 0 | 0 | 0 | 0 | 0 | 0 | 0 | 0 | 0 |
| 21 | "great job! :)" | 0 | 0 | 0 | 0 | 0 | 1 | 0 | 0 | 0 |
| 22 | N/A | 0 | 0 | 0 | 0 | 0 | 0 | 0 | 0 | 0 |
| 23 | "1 real lit game, minecraft 3" | 0 | 0 | 0 | 0 | 0 | 1 | 0 | 0 | 0 |
| 24 | "I like the approach to learn things through video games. It's easier to focus." | 0 | 0 | 0 | 1 | 0 | 0 | 0 | 0 | 0 |
| 25 | N/A | 0 | 0 | 0 | 0 | 0 | 0 | 0 | 0 | 0 |
| 26 | N/A | 0 | 0 | 0 | 0 | 0 | 0 | 0 | 0 | 0 |
| 27 | N/A | 0 | 0 | 0 | 0 | 0 | 0 | 0 | 0 | 0 |
| 28 | "Would be nice for teaching in schools" | 0 | 0 | 0 | 1 | 0 | 0 | 0 | 0 | 0 |
| 31 | "Can you lose at this game?" | 0 | 0 | 1 | 0 | 0 | 0 | 0 | 0 | 0 |
| 32 | "the game has the perfect length you're able to try out different decisions in each new game that you start which makes it easier to understand the value of microbiome diversity and also the effect of it" | 0 | 0 | 0 | 1 | 0 | 1 | 0 | 0 | 0 |
| 37 | N/A | 0 | 0 | 0 | 0 | 0 | 0 | 0 | 0 | 0 |
| 39 | N/A | 0 | 0 | 0 | 0 | 0 | 0 | 0 | 0 | 0 |
| | | **Expert group (*N* = 22)** | | | | | | | | |
| 5 | N/A | 0 | 0 | 0 | 0 | 0 | 0 | 0 | 0 | 0 |
| 7 | N/A | 0 | 0 | 0 | 0 | 0 | 0 | 0 | 0 | 0 |
| 8 | "some cut-szenes, videos, entertainment, user feedback would be great..." | 0 | 0 | 1 | 0 | 0 | 0 | 0 | 0 | 0 |
| 9 | N/A | 0 | 0 | 0 | 0 | 0 | 0 | 0 | 0 | 0 |
| 11 | N/A | 0 | 0 | 0 | 0 | 0 | 0 | 0 | 0 | 0 |

| | | | | | | | | | | |
|---|---|---|---|---|---|---|---|---|---|---|
| 12 | "Nice idea! Maybe get in touch with some schools" | 0 | 0 | 0 | 1 | 0 | 0 | 0 | 0 | 0 |
| 13 | N/A | 0 | 0 | 0 | 0 | 0 | 0 | 0 | 0 | 0 |
| 19 | N/A | 0 | 0 | 0 | 0 | 0 | 0 | 0 | 0 | 0 |
| 20 | "I couldn't hear any music. The full-screen mode cuts out some text and the arrows that allow you to move on from the book of decisions." | 0 | 0 | 0 | 0 | 0 | 0 | 0 | 1 | 0 |
| 21 | N/A | 0 | 0 | 0 | 0 | 0 | 0 | 0 | 0 | 0 |
| 22 | N/A | 0 | 0 | 0 | 0 | 0 | 0 | 0 | 0 | 0 |
| 23 | N/A | 0 | 0 | 0 | 0 | 0 | 0 | 0 | 0 | 0 |
| 24 | N/A | 0 | 0 | 0 | 0 | 0 | 0 | 0 | 0 | 0 |
| 27 | N/A | 0 | 0 | 0 | 0 | 0 | 0 | 0 | 0 | 0 |
| 28 | N/A | 0 | 0 | 0 | 0 | 0 | 0 | 0 | 0 | 0 |
| 30 | N/A | 0 | 0 | 0 | 0 | 0 | 0 | 0 | 0 | 0 |
| 31 | N/A | 0 | 0 | 0 | 0 | 0 | 0 | 0 | 0 | 0 |
| 34 | "Nope, the game is indeed achieving the goal!" | 0 | 0 | 0 | 0 | 0 | 1 | 0 | 0 | 0 |
| 40 | "the part with fetal microbiome is quite controversial according to Kennedy et al 2023 disproved https://www.nature.com/articles/s41586-022-05546-8. It would also be beneficial to include source like Walker & Hoyles 2023 https://www.nature.com/articles/s41564-023-01426-7. anyhow, a great work! I really enjoyed it!" | 0 | 0 | 0 | 0 | 0 | 0 | 0 | 0 | 1 |
| 41 | "I don't know if ethically it is possible to modify this, but personally I think looking at the citations made it feel less like a fun game and more like reading a scientific article. I understand why they are there, but just wanted to say my opinion, also thinking that maybe the game is directed to people that maybe are not so involved in science." | 0 | 0 | 1 | 0 | 0 | 0 | 0 | 0 | 0 |
| 43 | "look forward to the next level, if any." | 0 | 0 | 0 | 0 | 0 | 1 | 0 | 0 | 0 |
| 45 | N/A | 0 | 0 | 0 | 0 | 0 | 0 | 0 | 0 | 0 |

N/A: no answer given by the study participant
